# Directional biases in whole hand motion perception revealed by mid-air tactile stimulation

**DOI:** 10.1101/2020.04.23.058024

**Authors:** Marlou N Perquin, Mason Taylor, Jarred Lorusso, James Kolasinski

## Abstract

Human machine interfaces are increasingly designed to reduce our reliance on the dominantly used senses of vision and audition. Many emerging technologies are attempting to convey complex spatiotemporal information via tactile percepts shown to be effective in the visual domain, such as shape and motion. Despite the intuitive appeal of touch as a method of feedback, we do not know to what extent the hand can substitute for the retina in this way. Here we ask whether the tactile system can be used to perceive complex whole hand motion stimuli, and whether it exhibits the same kind of established perceptual biases as reported in the visual domain. Using ultrasound stimulation, we were able to project complex moving dot percepts onto the palm in mid-air, over 30cm above an emitter device. We generated dot kinetogram stimuli involving motion in three different directional axes (‘Horizontal’, ‘Vertical’, and ‘Oblique’) on the ventral surface of the hand. We found clear evidence that participants were able to discriminate tactile motion direction. Furthermore, there was a marked directional bias in motion perception: participants were better and more confident at discriminating motion in the vertical and horizontal axes of the hand, compared to those stimuli moving obliquely. This pattern directly mirrors the perceptional biases that have been robustly reported in the visual field, termed the ‘Oblique Effect’. These data show the existence of biases in motion perception that transcend sensory modality. Furthermore, we extend the Oblique Effect to a whole hand scale, using motion stimuli presented on the broad and relatively low acuity surface of the palm, away from the densely innervated and much studied fingertips. These findings also highlight targeted ultrasound stimulation as a versatile means by which to convey potentially complex spatial and temporal information without the need for a user to wear or touch a device. This ability is particularly attractive as a potential feedback mechanism for application in contact-free human machine interfaces.

## Introduction

As technology pervades almost every aspect of the built environment, the challenge of designing effective human machine interfaces (HMIs) increases [Proctor and Zandt, 2018]. In the modern era, vision in particular, and audition to a lesser extent, have dominated our interaction with technology. Screen-based interfaces are omnipresent, from self-service machines in shops and transport hubs, to the touch screen elements of cars, phones, and even home appliances. However, the attentional demands associated with complex HMIs risks sensory overload, meaning that the amount of incoming information is too high to be adequately perceived and acted upon [Woods et al., 2002]. Even in healthy young individuals, this could have severe consequences on performance in high-risk contexts, such as driving [Engström et al., 2005] or operating other complex control systems.

It has been argued that in specific circumstances humans can better perceive information conveyed across multiple distinct sensory modalities compared with the same volume of information communicated via a single modality (i.e. Multiple Resource Theory; [Wickens, 2008]). As such, the threshold for sensory overload may be increased by careful design of multisensory interfaces. This has prompted the development of new technologies that allow humans to perceive information beyond the dominant senses of vision and audition.

Tactile interfaces represent one of such growing technologies, and capitalise on our sense of mechanoreception (touch). This technology is not inherently new: braille is a clear example of a way in which the tactile domain can perform a perceptual function typically reliant on vision. More recently, tactile stimulation systems have shown promise as an alternative means of conveying information in HMIs, for example, in the context of driving and operating surgical robots [Meng and Spence, 2015, Okamura, 2009]. However, key technological shortcomings of tactile systems have included the limited ability to convey complex spatial information and the need to wear or touch an interface for a tactile signal to be conveyed.

A substantial advance in targeted ultrasound technology has overcome these limitations, making it possible to project complex tactile percepts projected onto the hand without any physical contact [Carter et al., 2013] (Figure 1). Mid-air touch stimulation uses an array of ultrasound emitters and infra-red hand tracking to deliver stimuli with a high spatial and temporal frequency, targeted to specific regions of the palmar surface of the hand, up to 80cm above an emitter device. Just as light is projected onto the retina for vision, ultrasound technology can project tactile scenes directly onto the hand. These can include defined points, lines, and shapes; both static and moving.

**Figure 1.**
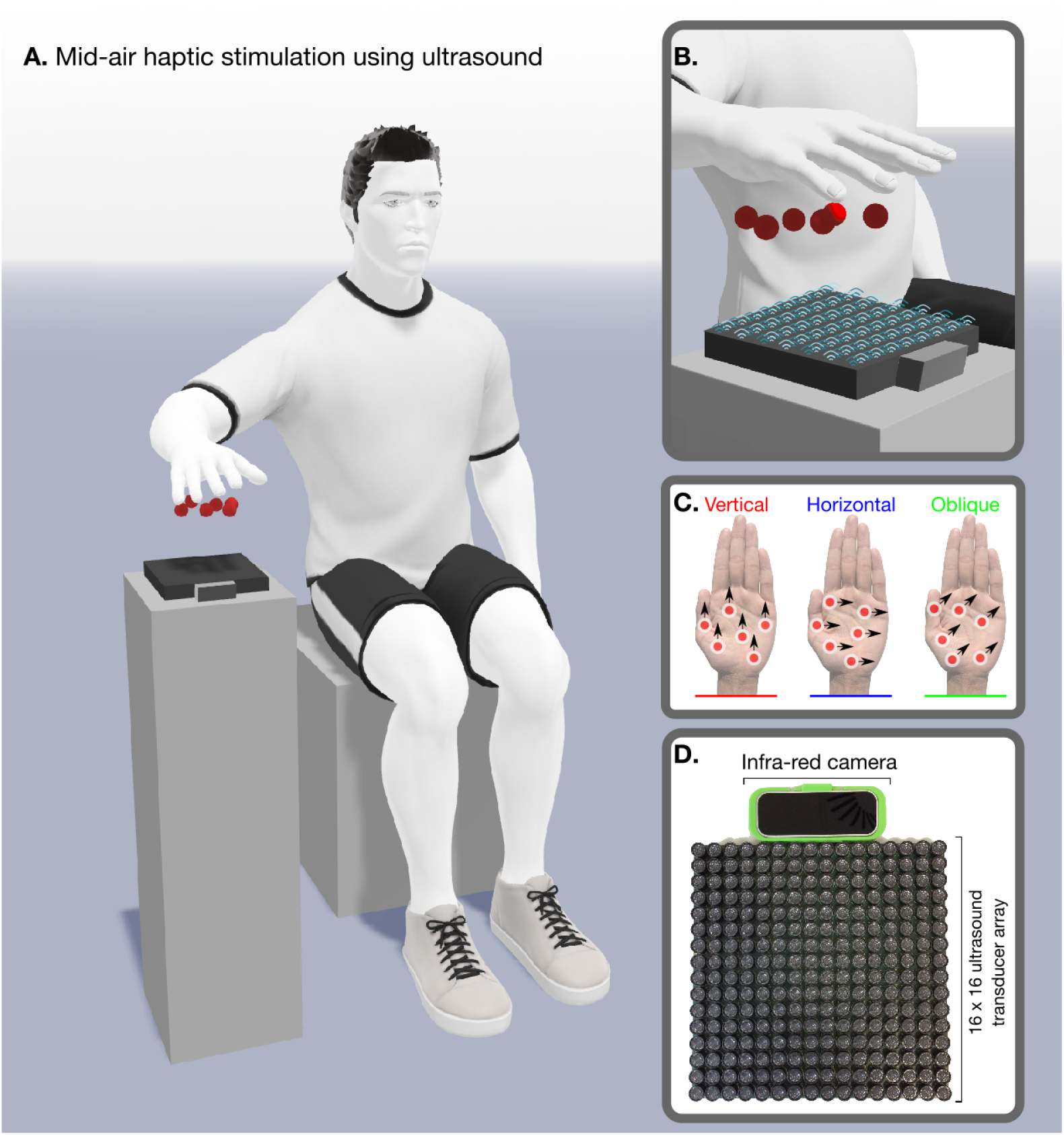
Overview of mid-air tactile experimental setup. (A) Participants were seated with their hands above an array of ultrasound actuators and a infra-red camera. (B) The combination of real time hand tracking and focused ultrasound can project discrete points onto the user’s unadorned hand [Carter et al., 2013]. (C) Users experienced a series of moving dot stimuli (Figure 2) in differing directions. (D) Stimuli were delivered using an Ultrahaptics device (UltraLeap, Bristol).

The development of advanced stimulation technology has arguably outstripped our understanding of human tactile perception. Of particular importance is the question of whether the hand can be used to perceive the relatively complex stimuli that the technology can emit. Although the spatial and temporal features of such whole hand tactile stimuli prompt obvious parallels with visual stimuli (shapes, lines, motion), touch is a distinct sensory domain. The potential application of such technology to HMIs thus currently hinges on the assumption that humans can perceive the spatial and temporal features whole hand tactile stimuli in the same way we would perceive equivalent visual stimuli with our eyes. Empirical evidence is lacking, because studies of human touch sensation on the hand have been dominated by investigation of the fine-grain perceptual abilities of small regions of high tactile acuity at the fingertips [Mountcastle, 2005], rather than the ability to perceive tactile scenes projected across entire palm.

If the hand can indeed be used to perceive such complex stimuli similar to vision, one may also ask if it exhibits the same kinds of well-documented perceptual biases reported extensively in the visual domain. This is of importance, as the presence of such common biases across the visual and tactile modalities would further support the dominant notion of integrative multisensory processing in the human brain [Murray et al., 2016]. A clear understanding of these biases in whole hand tactile function is essential to the development of HMIs that work in synergy with human perceptual abilities.

In this work we address these questions by using novel focused ultrasound stimuli to translate a classic visual dot kinetogram stimulus to tactile domain. We utilise the visual oblique effect [Appelle, 1972], which refers to a well-established phenomenon that perception of motion or orientation in horizontal and vertical axes is superior to that in intermediate oblique axes (e.g., [Ball and Sekuler, 1980, 1987]). Specifically, we ask (1) whether the human hand can be used to accurately and confidently perceive the complex dot motion stimuli in the tactile domain, and (2), whether the oblique effect manifests in perception of tactile motion stimuli presented in different orientations on the palm.

## Materials and Methods

### Participants

Fourty-five participants (29 female, age range: 19-40, Mean_age_ = 24.2) were tested in a two-day experiment. Participants were right-handed, with no self-reported touch deficits in their hands or upper limbs, and were paid 30 pounds in total for participation. The study was approved by the local ethics committee (Cardiff School of Psychology Research Ethics Committee: EC.18.06.12.5311R). Three participants were excluded from analysis: the first responded almost exclusively with one response key; the other two were excluded because of excessive movement, which led to difficulty in tracking both their hands as well as their eye movements.

### Materials

The experiment was generated using PsychoPy 3.2 [Peirce, 2007, 2009, Peirce et al., 2019] and Visual Studio 16 (Microsoft, Redmond, US), run on a Viglen Vig800S computer (Viglen Ltd, Hertfordshire, UK). The tactile stimuli were generated from an UltraHaptics UHEV1 Array (UltraLeap, Bristol, UK), which was attached with a Leap Motion camera for continuous hand-tracking. The visual instructions and elements of the experiment (Figure 2) were displayed on an ASUS VG248QE monitor (resolution: 1920×1080; refresh rate: 144 Hz; AsusTek Computers Inc, Taipei, Taiwan). Participants’ eyes were tracked with a LiveTrack Lightning Eye Tracker (Cambridge Research Systems Ltd, Kent, UK). Responses were recorded with a NAtA Technologies Response Box (NAtA Technologies, Coquitlam, Canada). Hand and body temperature were recorded with an NC200 Non Contact Forehead Thermometer (Medisave UK Ltd, Weymouth, UK). 3D visualisations of the experimental paradigm were generated using MagicPoser (Wombat Studio, Inc., Santa Clara, California, US).

**Figure 2.**
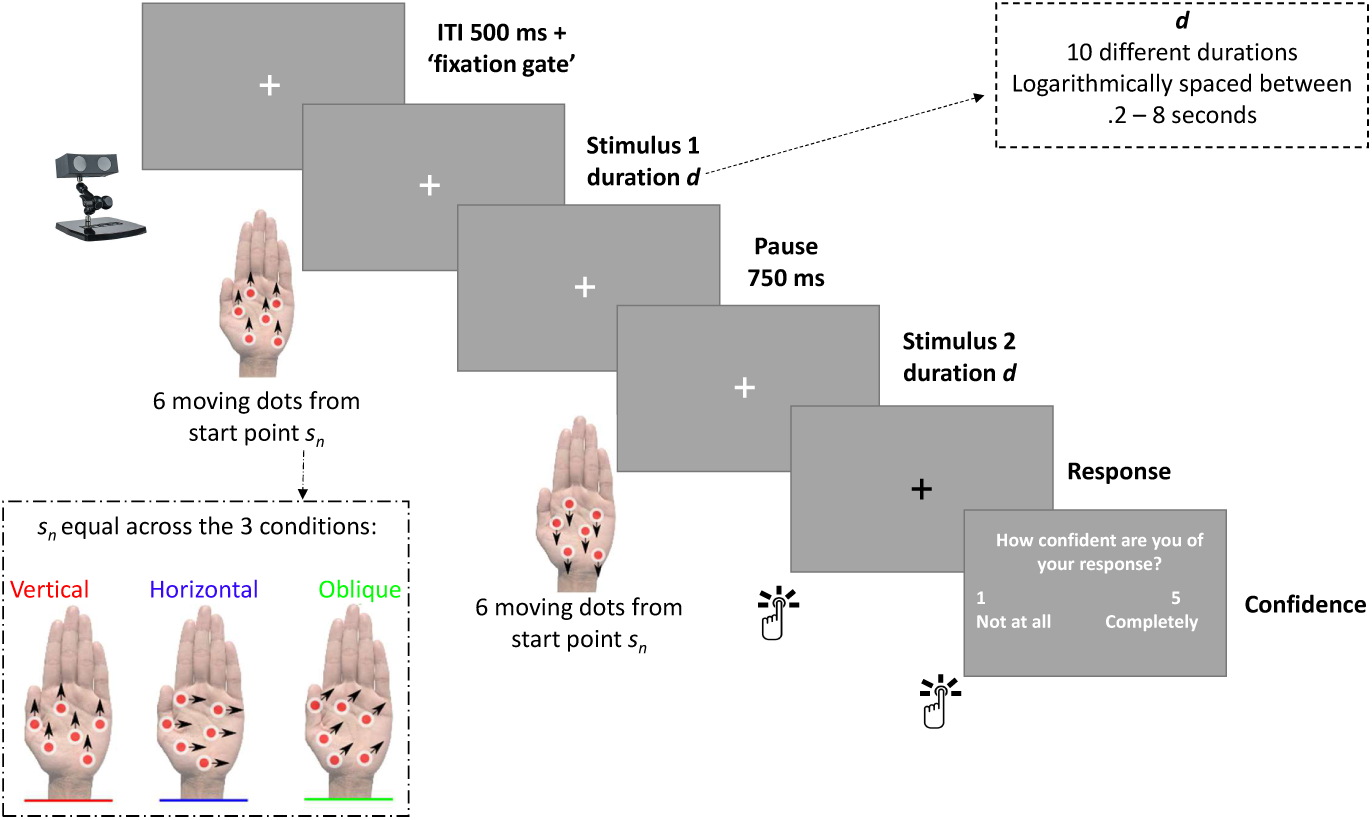
Overview of a single experimental trial presenting midair tactile stimuli in a two-interval-forced-choice task. In the training block, each trial started with a white fixation cross, presented for 500 ms. The first stimulus was then delivered for 1 second, followed by a 750 ms pause, and then the second stimulus was delivered for 1 s. The motion stimuli consisted of six tactile dots generated by the Ultrahaptics array starting from *s*_n_, with *s* being a randomly-assigned start point at trial *n*. Note that *s*_n_ was kept equal over the three conditions (i.e. for one participant, the start position of the six random dots on trial *n* were the same in each condition). In addition, motion was limited such that the moving dots would not extend beyond the palmar area (meaning participants would always be able to feel motion for the full stimulus duration). Any dots extending beyond the motion area were re-generated at a pseudorandom position in the motion area. After stimulus presentation, the fixation cross turned black, prompting a response from the participant. After a correct response, the cross turned green, and after an incorrect response, the cross turned red. Participants completed six trials for each condition (horizontal, vertical, and oblique), with a self-passed break in between each condition. Condition order was counterbalanced across participants.

### Design

Each participant took part in a two-interval-forced-choice task, in which they were instructed to discriminate tactile motion direction. This was undertaken in three different conditions (Figure 2): 1) horizontal (with stimuli moving along the 90-270^°^ axis, along the medial-lateral axis of the palm; one stimulus would move to the left, and the other stimulus would move to the right), 2) vertical (0-180^°^ axis, along the proximal-distal axis of the palm), and 3) oblique (45-225^°^ axis). After the presentation of the two stimuli in a given axis, participants judged which of the two stimuli respectively moved: 1) rightwards, 2) downwards, and 3) oblique downwards.

All stimuli consisted of 6 tactile dots (8.5mm diameter), moving coherently at a speed of 4cm/second. Dots were selected as they do not provide any other potential motion cues, such as shape and orientation. The area on the palm of the hand in which the motion occurred extended across the full medial to lateral extent of the palmar surface. The proximal-distal extent of the palmar motion area was equal to the medial to lateral width of the palmar surface extending from the heel of the hand to the proximal aspect of the fingers.

Previous pilot studies using Ultrahaptics [Korres and Eid, 2016, Rutten et al., 2019] have selected only very long stimulus durations, respectively lasting 9 and 30 seconds. As the consequences of such long durations are still unknown, we presented stimuli across a broad range of 10 different durations *d*, which were logarithmically spaced between 200 ms and 8 seconds, allowing us to investigate the potential for accurate perception across a wide variety of exposures.

### Procedure

Participants visited the lab for two sessions, each 1-1.5 hour in length. In the first session, they completed the Edinburgh Handedness Inventory [Oldfield, 1971] to verify their right-handedness. Next, they were seated in a chin-rest, 55 cm from the screen. Their right arm was immobilised on an arm-rest with velcro straps to eliminate motion, with their hand 38 cm above the Ultrahaptics device. To block any auditory cues from the presented stimuli, participants were given 35 dB ear plugs as well as a pair of 27.6 dB ear defenders. During the training period participants wore the ear defenders without ear plugs to allow for verbal discussion and questions.

Participants first performed a training block. Next, they were taken out of the arm-rest, to ask any questions and to fit ear plugs. After the training, the eye tracker was calibrated using a 9-point paradigm. Participants subsequently performed two experimental blocks of each condition. In the second session, participants performed three more experimental blocks per condition.

The experimental blocks, which comprised the majority of the testing, mirrored the training blocks aside from three key differences. Firstly, each trial included a ‘fixation gate’ - a gate for which participants had to continuously fixate for ∼175 ms (25 frames) before the trial would begin - presented after the initial fixation dot of 500 ms. Secondly, participants were not given feedback after their responses, but instead were asked to rate the confidence in their answer on a scale from 1: “not at all confident”) to 5: “completely confident” (Figure 2). Finally, the stimulus duration was varied across trials at intervals logarithmically spaced between 200 ms and 8 seconds.

Each experimental block consisted of 40 trials, with 4 times each of the 10 durations, randomly presented throughout block (but see section: Technical issues); each block comprised only one condition. There was a self-paced break after every 10 trials, and a longer break between each condition - this large number of breaks was included to minimise discomfort and to reduce participant motion during the trials.

During pilot studies preceding the current experiment, large differences in performance and confidence were already apparent between participants. Aiming to explain these differences, we measured a number of candidate variables that may affect tactile performance. At the beginning of both sessions, body temperature (measured on the forehead and the right hand), hand size (measuring from the wrist to the tip of the middle finger), and (middle) finger length were also measured. We also considered task effects on performance, namely (between-subject) condition order and (within-subject) time on task (i.e., block number).

### Technical issues

A common technical issue we experienced was the Ultrahaptics emitter crashing within a trial - possibly because the device is not specifically made for experimental purposes, in which it is necessary to run a large number of trials in quick succession. To limit the amount of these crashes, we reconnected the device after every 40-trial block. However, we were not able to prevent the crashes altogether, and still had 157 crashes over all sessions combined (3.7 crashes per participant on average, SD = 2.8, range 0 to 12). Wherever possible, participants performed extra trials, aiming to achieve a *minimum* of 200 trials in each condition (total trial mean across 3 conditions = 614, SD = 25, range: 560 to 660). Number of trials did not vary systematically between condition, BF_01_ = 12.5.

### Data preparation and analysis

Training trials were excluded from all analyses. Analyses were conducted in Matlab 2019a [MATLAB, 2019] and JASP [JASP Team, 2019]. As this is the first study to systematically investigate the feasibility of tactile mid-air perception, we chose to use Bayesian statistics - allowing for assessment of both the alternative and the null hypothesis. Bayesian statistics were estimated using equal prior distributions and 10,000 iterations for Monte Carlo simulations.

### Performance

#### Feasibility of tactile mid-air perception

Participant means for performance (% correct) were calculated for each of the three conditions separately, as well as combined over all conditions. Bayesian one-sided one-sample t-tests were conducted on the group distributions, to test if they were higher than chance level (50 % correct).

#### Testing within-subject effects

Participant means were calculated separately for each condition and duration. A Bayesian 3×10 Repeated Measures (RM) ANOVA was conducted, with condition and duration as independent factors.

### Confidence

#### AROC: Quantifying meta-cognitive ability

To assess participants’ subjective experiences of the tactile stimuli, confidence ratings were measured after every trial. However, raw confidence ratings cannot quantify how *accurate* the participant is at judging their own performance (i.e., high ratings for correct responses, and low ratings for incorrect responses). To assess such ‘meta-cognitive ability’, we also estimated the type-II area under the receiver-operating curve (AROC; [Fleming and Lau, 2014]; [Fleming et al., 2010]). Just as in behaviour, one could distinguish *hits* (in this case, a high confidence rating when the response is correct) from *misses* (a high confidence rating when the response is incorrect). To make such a distinction, one needs a criterion to determine if the rating is ‘high’ or ‘low’. To estimate the ROC, the proportion of hits can be plotted against the proportion of misses along all possible criteria (that is, the proportion calculated under low = 1 and high = 2-5, the proportion calculated under low = 1-2 and high = 3-5, and so on). Just as with typical ROCs, the area under the curve can then be quantified, giving the AROC measure. A key benefit of the type-II AROC is that it does not assume the confidence ratings follow a normal distribution - an assumption that is not met in the current data.

Chance level of the AROC measure is indicated by a value of 0.5. To assess whether the current values exceeded this level, Bayesian one-sided one-sample t-tests were conducted on the group distributions of AROC values for all three conditions plus combined.

#### Testing within-subject effects

Mean confidence and AROC were calculated per participant separately for each condition and duration. Two Bayesian 3×10 Repeated Measures (RM) ANOVAs were conducted using condition and duration as independent factors.

### Examining inter-individual differences and task-effects

#### Between-subject correlations

Participants’ age, hand size, and finger size were linearly correlated between individuals to mean accuracy, confidence, and AROC (looking at combined, horizontal, vertical, and oblique trials over the entire session) - resulting in distributions of (3×3×4) 36 correlation coefficients plus accompanying Bayes Factors. Bayesian ANOVAs were calculated on the same outcome measures, with condition order as independent variable - giving 12 additional Bayes Factors.

#### Within-subject correlations

To assess within-subject factors, accuracy, confidence, and AROC were calculated for each of the five blocks, independently for combined, horizontal, vertical, and oblique trials. For each participant, these means were correlated to: 1) hand temperature prior to each block, 2) hand temperature before each block, corrected for body temperature, and 3) block number. This resulted in (3×3×4) 36 correlation coefficients per participant. We tested whether each of these coefficients were statistically different from zero on the group level, using Bayesian one sample t-tests.

#### Interpretation Bayes Factors

BF_10_ represents the likelihood of the current data under the alternative (e.g., effect of condition) over the null hypothesis (e.g., no effect of condition). It is a continuous measure of evidence that can take any value between zero to infinity. Note that the evidence for the null over the alternative hypothesis (BF_01_) is equal to the inverse of BF_10_. BF_10_ values above 1 indicate more evidence for the alternative hypothesis, while values under 1 indicate more evidence for the null-hypothesis - though as a rough rule of thumb, BF_10_ between 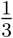 and 3 are typically interpreted as ‘indeterminate evidence’.

Bayesian RM ANOVA is a form of model comparison - assessing how much more likely the data is under the statistically-best model as compared to under each of the other models. The output provides an ‘Analysis of Effects’, with BF_inclusion_ reflecting the average over all the models which include that factor; this is therefore the most comparable to ‘classic’ RM ANOVA within-subject effects. The Bayesian RM ANOVA also provides Bayes Factors to compare the models directly. In our current analyses, the model comparisons and the analyses of effects led to the exact same conclusions in each instance. Therefore, we chose to only report the BF_inclusion_. The Bayes Factors for model comparison, as well as all other analyses, can be found on the annotated .jasp files on OSF.

## Results

### Performance

#### Discrimination of complex tactile percepts exceeds chance

On the group level, there was extreme evidence that performance was statistically above chance in all conditions (Figure 3A), indicating that participants were able to distinguish the direction of complex tactile motion perceptions delivered across the palm. The highest evidence for performance above chance was observed in the vertical condition and the lowest was observed in the oblique condition. Figure 3A shows the accuracy mean for each participant for both average performance and separately over the three conditions, with accompanying BF_10_ above. Sequential analyses reveal the trajectory of evidence accumulation across participant recruitment, reflecting the change in BF_10_ as the sample size increased. Average overall accuracy across all stimulus durations was not very high (overall group mean = 60.6%) and between-subjects variance was high.

**Figure 3.**
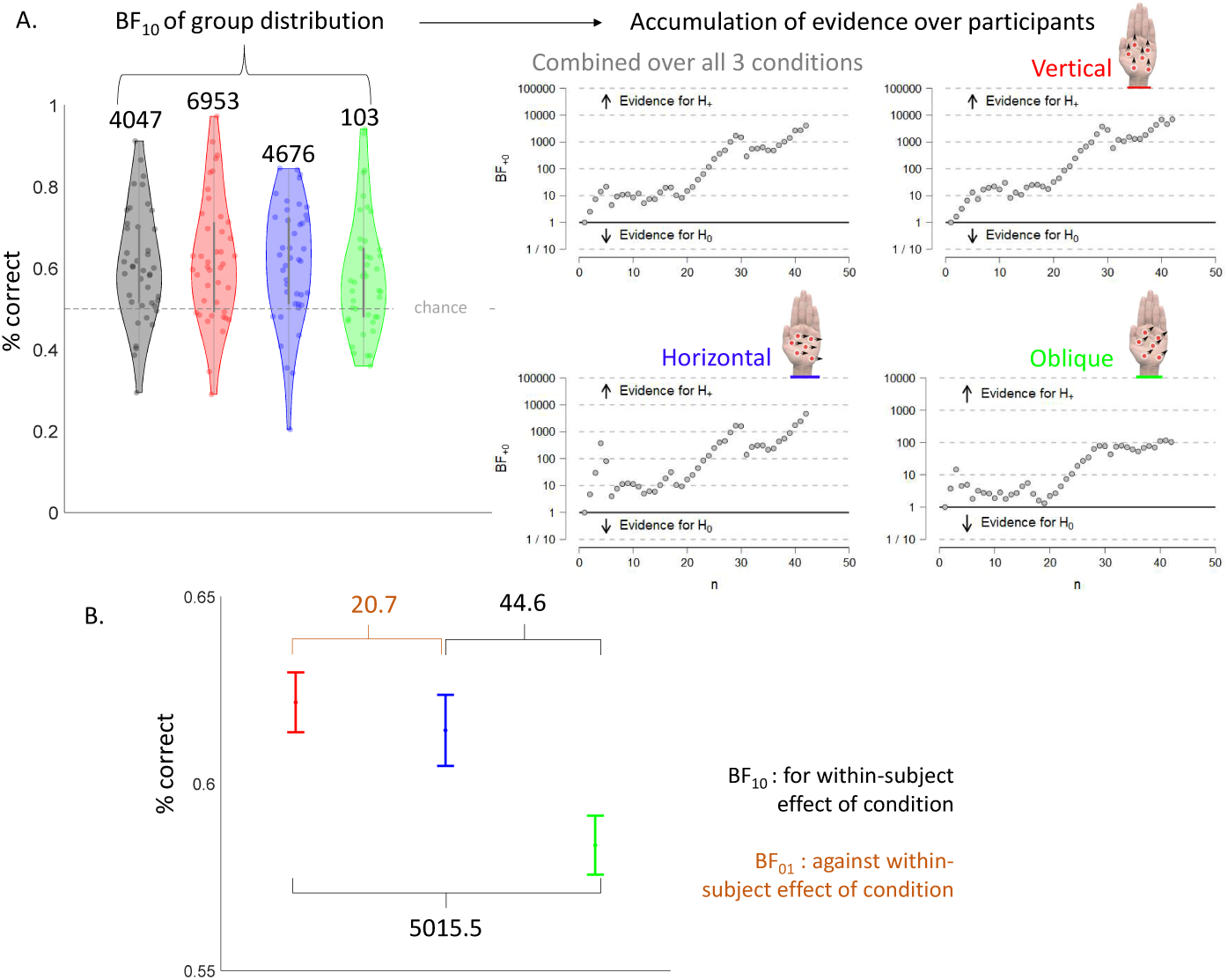
Evidence of an oblique effect in whole hand tactile motion perception. Performance on the experimental two-interval-forced-choice task for the three conditions combined (grey) as well as separate for vertical (red), horizontal (blue), and oblique (green) tactile motion stimuli presented on the palm. A. Group distributions of % correct (left), with each dot in a distribution showing the performance of one participant, with the accompanying BF_10_ from the one-sample t-tests presented above. The BF_10_ show extreme evidence that % correct exceeds chance level. The right panel shows the change in BF_10_ as a function of participant recruitment, reflecting the accumulation of evidence as the sample size increased. B. Plot of the group mean of each condition with the within-subject error bars - reflecting the within-subject differences across the three conditions - with the BF_10_ of the post-hoc tests above. Post-hoc tests indicate a clear existence of an “Oblique” effect in the data, such that participants performed statistically better in perceptual discrimination in the horizontal and vertical axes compared with the oblique axis.

#### Clear evidence of an oblique effect in tactile motion perception

Figure 3B shows the within-subject differences between the three conditions, with accompanying statistics in Table 1. The BF_inclusion_ indicate that there is extreme evidence for an effect of condition only.

**Table 1.** Overview of the BF_inclusion_ for the three independent factors - condition, duration, and their interaction - resulting from the three RM ANOVAs conducted on performance (% correct), confidence rating, and metacognitive ability (AROC). BF_inclusion_ that indicate evidence in favour of an effect (> 3) are shown in green; BF_inclusion_ that indicate evidence against an effect 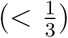 are shown in red.

| Factor | % Correct | Confidence | AROC |
| --- | --- | --- | --- |
| condition | 1056 | 6.6e+7 | 0.006 |
| duration | 7.0e-5 | 5.3e+7 | 3.6e-5 |
| conditon*duration | 1.6e-7 | 1.99e-4 | 1.98e-8 |

Post-hoc tests conducted on condition (Bayes Factors shown in Figure 3B) showed that accuracy was lower in the oblique compared to the vertical and horizontal condition, with no difference between the horizontal and vertical conditions. The results indicate that participants performed significantly better in perceptual discrimination of tactile motion presented along the horizontal and vertical axes compared with the oblique axis, consistent with the notion of an oblique effect from the visual literature.

#### Confidence in tactile perception also shows oblique effect

Mean confidence over participants and conditions on a 5-point scale was 3.1 (SD = .14), indicating that on average participants felt neutral about the accuracy of their responses: neither very confident nor very unconfident. Mean AROC was 0.57 (SD = 0.10), with Bayesian one-sided one sample t-tests showing the distributions were higher than 0.5 (BF_10_ = 465, 80, 3337, and 26 for combined, vertical, horizontal, and oblique motion conditions respectively). Figure 4A shows the break-down of confidence and AROC over the three conditions and ten durations.

**Figure 4.**
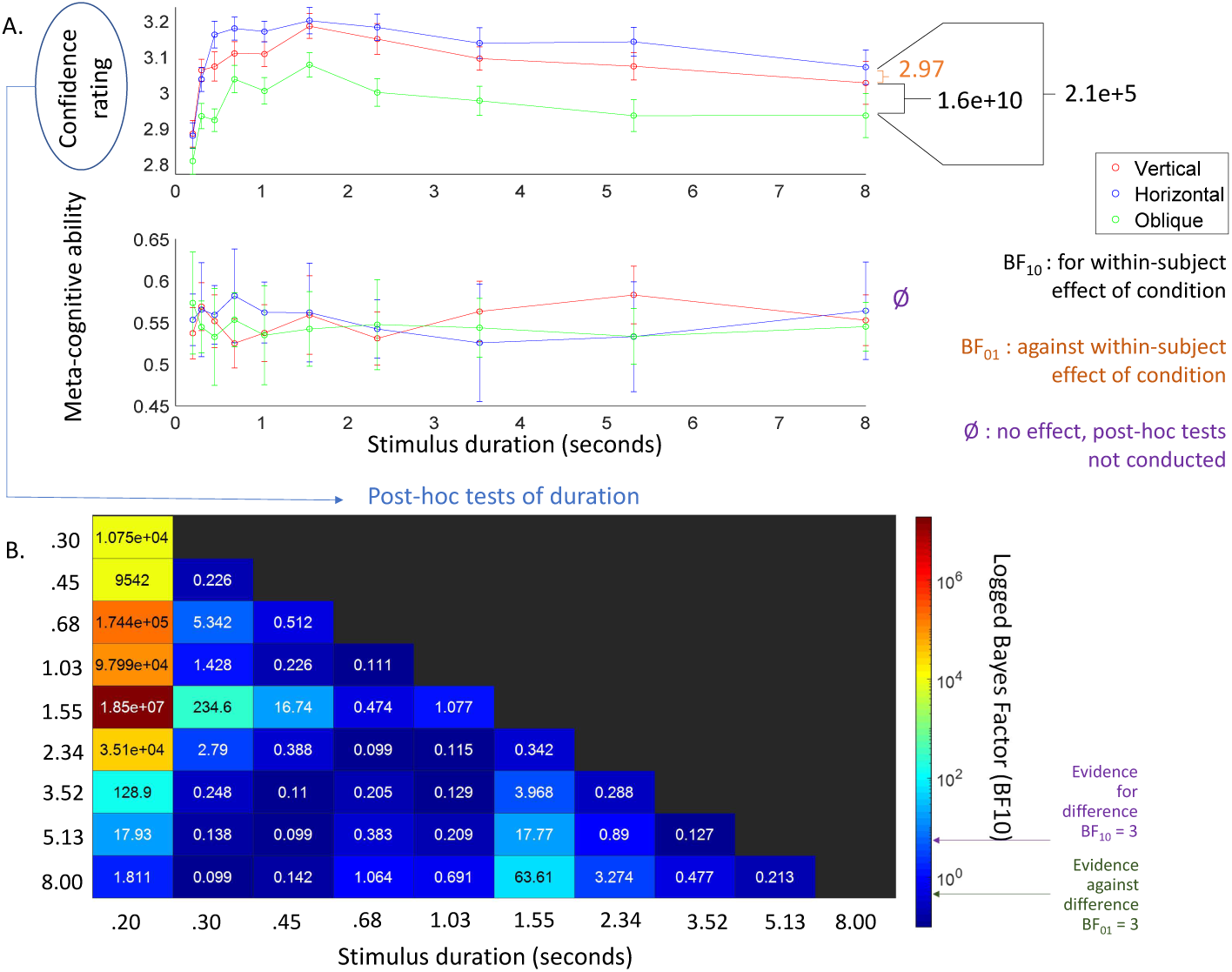
Overview of results of the meta-cognitive measures. A. Mean confidence rating (top panel) and mean meta-cognitive ability (AROC; bottom panel) for the vertical (red), horizontal (blue), and oblique (green) condition over the ten different stimuli durations (logarithmically spaced between 200 ms and 8 s). Error bars indicate the within-subject error across conditions. There was a main effect of condition and duration on confidence rating, but not on meta-cognitive ability. On the right, the BF_10_ from the post-hoc tests on condition are shown. Again, there is a clear oblique effect, with confidence in the oblique condition being worse than in the horizontal and vertical condition. B. The BF_10_ from the post-hoc tests on duration. Dark blue colours indicate more evidence for the null-hypothesis, while lighter blue to red colours indicate gradual higher evidence for the alternative hypothesis. Overall, confidence is lowest in the shortest and in the longest durations.

**Figure 5.**
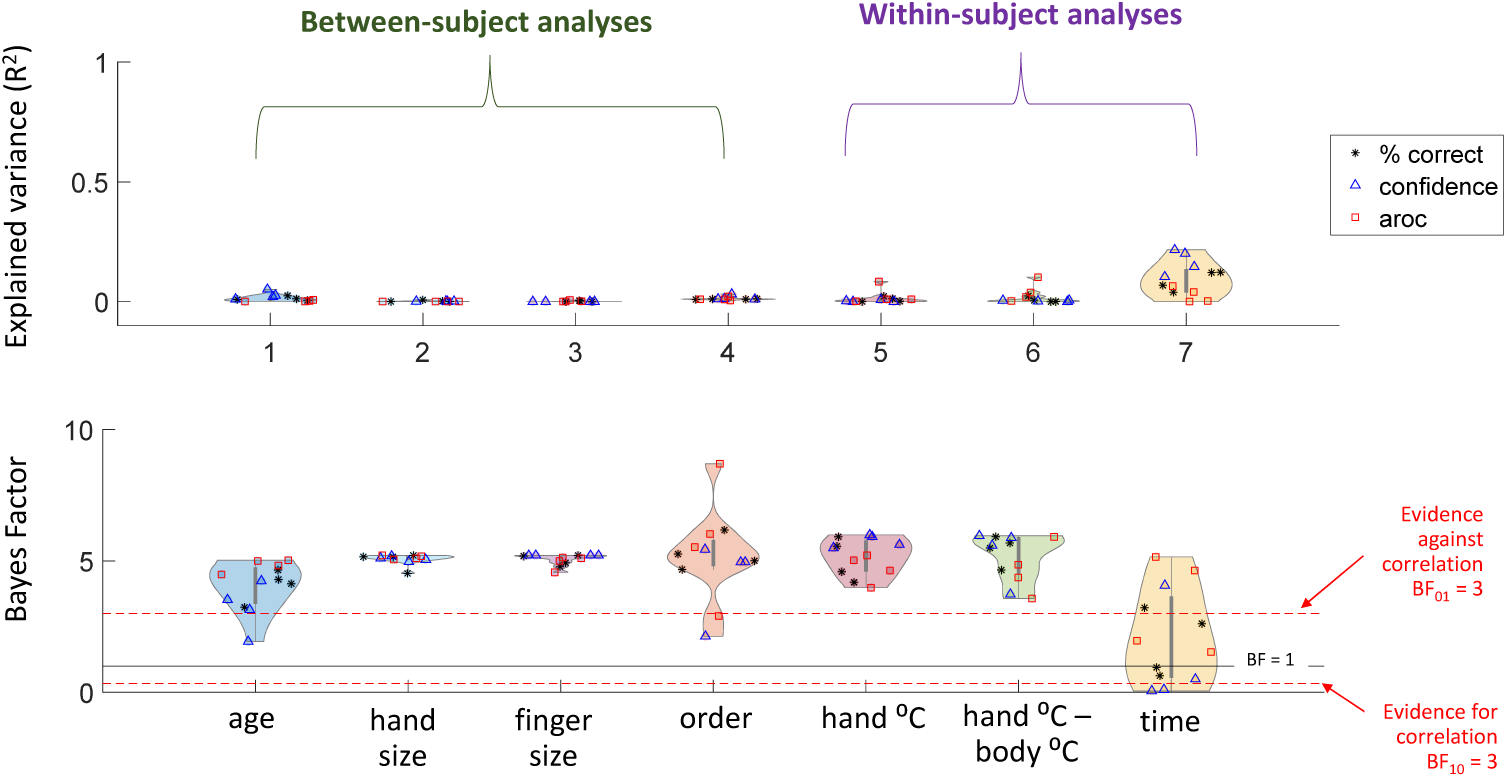
Distributions of the explained variance (R^2^; top panel) and accompanying Bayes Factors (bottom panel) of the correlation analyses and ANOVA of individual differences and task effects. Analyses were conducted on performance (star), confidence ratings (triangle), and AROC (square). Analyses are separated between those conducted on the between-subject (left) and on the within-subject level (right). In the bottom panel, values above the upper red line indicate more evidence for the null-hypothesis (BF_01_ > 3). Values between the two red lines are typically interpreted as indeterminate, while evidence below the red line 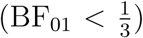 indicate evidence for the presence of a correlation/effect. Overall, we find evidence against systematic individual differences and task effects for age, hand size, finger size, order, and hand temperature. The evidence for effects of time remain largely indeterminate.

#### Stimulus duration and motion orientation affect confidence in tactile perception

There was extreme evidence for an effect of condition and of duration on participant confidence, but extreme evidence against an interaction-effect. Similarly to performance, post-hoc tests for condition (Figure 4A) showed extreme evidence that participants were less confident in the oblique compared to the horizontal and vertical condition, with moderate evidence against a difference between horizontal and vertical. Again, these data suggest that participants were significantly less confident in their perceptual judgements on the oblique axis.

Due to the large number (35) of post-hoc tests for duration, the logged BF*10* are presented as a heatmap in Figure 4B. Overall, the effect of duration on confidence rating assumed an inverted U-shape: confidence is lowest in the very short and the very long tactile stimulus durations - with the highest confidence ratings reported for durations between 680 to 2430 ms.

In contrast, there was extreme evidence against effects of condition, duration, and their interaction on the measure of meta-cognitive ability (AROC) showed no effect of condition. This suggests that participants’ reduced confidence ratings in the oblique condition do not reflect a decline in their sensitivity, but rather match their actual lower performance.

#### Examining inter-individual differences and task-effects on tactile perception

To systematically assess the large number of between- and within-subject analyses, the BF_01_ for each analysis is plotted in violin plots (Figure 4 - bottom panel), with one distribution for each variable. Distributions shifted above the top red line show evidence against correlations (or against an effect, for ‘order’). The accompanying explained variance (R^2^) is shown in the top panel. Note that for the within-subject analyses, R^2^ reflects the median of the group distribution.

Neither performance, confidence, or AROC correlated with any of the between-subject factors (age, hand size, and finger size). Likewise, none of the outcome measures were affected by condition order. Within-subject fluctuations in performance, confidence, or AROC were not caused by fluctuations in hand temperature - neither ‘raw’ or normalised by body temperature. Explained variance for all these six variables centered around 0%.

BF_01_ for within-subject correlations between the outcome measures and time were largely indeterminate. Some of the confidence ratings were positively correlated with time - indicating that participants felt more confident in performance as they got more experience. This was, however, not mirrored in their objective performance.

## Discussion

Here we investigated the ability of people to perceive complex whole hand tactile motion stimuli generated using cutting-edge mid-air ultrasound technology. On a fundamental level, we found that participants could discriminate direction above chance level across all motion axes under study, despite no physical contact between the hand and the stimulator. Furthermore, we report evidence of a clear anisotropy in the perception of tactile motion across the palmar surface of the hand. Specifically, performance was poorest for motion discrimination in the oblique axis compared to the horizontal and vertical axes (Figure 2). The observed pattern was mirrored in measures of subjective confidence: people felt least certain in their motion discrimination judgements when the stimuli were moving in the oblique axis. This finding extends the classic ‘Oblique effect’ [Appelle, 1972] reported in visual motion into the tactile system. By translating the classic studies of motion dot kinetogram from visual to tactile domain, we have provided further clear evidence of commonalities in perceptual biases that transcend sensory modality.

Our results raise new questions regarding the perception of complex tactile percepts that can be projected onto the palm. From a mechanistic perspective: what are the underlying shared processes in the brain that confer common biases in motion perception across differing sensory modalities? From an applications perspective, how can the perceptual predilections of the human brain be used design effective feedback for touch-free HMIs using mid-air stimulation?

### Anisotropy in tactile perception

Anisotropy in tactile perception of orientation has been reported widely on the fingertips. There is with some disagreement regarding the specific axes in which acuity for orientation is highest on the fingertips. Some studies have reported enhanced perception of static tactile grating stimuli when they are oriented in the proximal-distal axis, parallel to the papillary ridges [Schneider et al., 1986, Essock et al., 1992, Wheat and Goodwin, 2000, Vega-Bermudez and Johnson, 2004], however others have reported enhancement in the medial-lateral axis in addition [Lechelt, 1988, 1992], without enhancement in the vertical axis [Bensmaia et al., 2008], or even isotropic perception across all orientations [Craig, 1999]. Few studies have considered tactile motion anisotropy. In a study of fingertip motion using a braille pin mounted on a trackpad, evidence for superior perceptual abilities was again reported only in the vertical axis of the fingertip [Keyson and Houtsma, 1995]. Reports of direction-dependent perceptual acuity of tactile orientation and motion stimuli have commonly attributed these to the orientation of the skin at the fingertip and differential sensitivity around the tip of the nail.

Here we report an oblique effect on an entirely different scale, with stimuli that extend across the palm of the hand. Using contact free ultrasound methods, we were able to deliver stimuli closely analogous to the random dot kinetograms common in the visual literature. We observe clear evidence of relatively enhanced motion perception in the vertical and horizontal directions aligned with the proximal-distal and medial-lateral axes of the hand. The observation of an oblique effect on this scale is striking, and shows clear distinctions from the more mixed evidence reported by experiments delivering fine-grain stimuli over limited spatial areas at the fingertip. The palm has a much lower receptor density and lower tactile acuity than the fingertips [Johansson and Vallbo, 1979, Mancini et al., 2014], and the cortical representation is correspondingly much smaller [Mountcastle, 2005].

A recent study investigated size perception across the hand, including stimuli along the same three axes as were applied here [Fiori and Longo, 2018]. The orientational biases in size perception appeared quite distinct from the evidence of an oblique effect presented herein. While we observed evidence of enhanced motion perception in both the vertical and horizontal axes, this work showed that judgements of size were most accurate when stimuli were presented in the horizontal (medial-lateral) axis and least accurate for stimuli presented in the vertical axis. The authors related this pattern of results to a perceptual stretch model, wherein perceived distance varies sinusoidally as a function of stimulus orientation. Increasing stretch increases the magnitude of the sinusoid, magnifying perceptual biases of size in stimuli shifted away from the horizontal axis. A meta-analysis of similar size perception studies on the palm concluded that an anisotropy exists, such that distances in this axis are perceived as around 10% larger than those in the vertical axis [Longo, 2020]. Here we extend on this very recent work to demonstrate that the glabrous skin of the palm, which was previously not thought to show anisotropies in tactile size perception[Longo and Haggard, 2011], also shows a clear pattern of perceptual anisotropy in motion perception.

### Neural mechanisms of tactile motion perception

The oblique effect generally appears to be driven by both lower-level *Class-1* and higher level *Class-2* mechanisms in the brain. In the visual domain, *Class-1* mechanisms involve the presence of fewer neurons tuned to oblique orientations in primary visual cortex (V1) compared to those responsive to vertical and horizontal orientations [Essock, 1980, Li et al., 2003], while *Class-2* mechanisms involve higher-level processing, such as memory and learning effects [Essock, 1980]. It is not possible to dissociate *Class-1* and *Class-2* mechanisms of tactile perception from the present design. However, the origins of directional biases in the tactile representations can perhaps be linked to long-term patterns of sensory inputs to the system. Recent work used arrays of up to 30 miniature acceleratometers to measure the patterns of cutaneous vibrations that pass through human hands during single finger, multi-finger, and grasping motions [Shao et al., 2016]. This data revealed clear evidence of gradients of vibrational intensity induced sequentially by each movement, which show broad alignment with the cardinal vertical and horizontal axes of the hands. Given the frequency with which we use such movements to interact with the world around us, it seems conceivable that the combination of the anatomy of hand movement, combined with the experience of stereotyped vibrational inputs, shapes the neural tactile representations around the cardinal axes.

The known neural mechanisms of tactile motion perception and its commonalities with primate visual motion processing have been well outlined in two recent reviews [Pei and Bensmaia, 2014, Pack and Bensmaia, 2015]. Evidence from primate electrophysiology suggests that Brodmann Area 1 (BA1) in the postcentral gyrus integrates amplitude, direction, and speed information from primary cortical neurons to yield tuning to specific motion directions in relatively larger receptive fields than those observed in other regions of primary somatosensory cortex (S1) [Gardner, 1988, Pei et al., 2010, 2011]. In this sense, BA1 seems to subserve a similar function to the middle temporal (MT)/V5 complex in visual motion processing [Pack and Bensmaia, 2015]. Evidence from human studies suggests that tactile motion also elicits more widespread activity in anterior intraparietal and inferior parietal areas [Kitada et al., 2003], as well as activation of an area of MT distinct from that implicated in visual motion [Summers et al., 2009, Wacker et al., 2011, Amemiya et al., 2017]; however the latter may be an epiphenomenon of visual imagery [Lacey and Sathian, 2011]. Directional biases in tactile motion perception appear to be independent of visual input, as tactile perceptual anisotropy has been observed previously in blind individuals [Lechelt, 1988].

### Applications of tactile stimuli in touch-free human-machine interfaces

The design of tactile stimuli is still in its infancy compared to other areas of HMI. As tactile technologies advance, so to do the complexity and sophistication of the stimulation methods possible [Schneider et al., 2017]. The risk of such rapid advances is that they outpace our understanding of human perception and develop based on the notion that features robustly perceived in one sense can also be perceived in another.

The tactile stimuli under study in this experiment were purposely designed to uncover relative differences across the three motion axes purely in the context of motion, and to avoid a ceiling effect in any one condition, hence their relatively small size (8.5mm diameter). When comparing receptor densities across the palm and retina, it is unsurprising that overall performance in the perception of these tactile motion ability remained relatively low. Making calculations based on reference densities of rapidly adapting receptor in the palm (0.92 receptors/*cm*^2^) [Johansson and Vallbo, 1979], our 0.85cm diameter tactile dots would excite fewer than one receptor per frame of movement. In contrast, using reference angular cone density data from the retina, an equivalent visual dot viewed at an equivalent distance (38cm) would excite around 180 cones in the fovea (assuming angular cone density of 180, 000*/degree*^2^) [Wang et al., 2019] or on average across the entire retinal surface, around 4-5 cones per frame of movement (assuming average angular cone density of 350*/degree*^2^) [Curcio et al., 1990]. On the basis of these figures, it is unsurprising that the stimuli were challenging to perceive, with an overall mean 61%, while an equivalent visual task would prove simple. Clearly larger non-overlapping dots would activate a large number of peripheral receptors and potentially enhance accuracy using a stimulus that remains purely motion-based.

By studying these challenging stimuli in isolation of complementary features such as orientation or shape, which might implicitly aid motion discrimination judgments, we are able to isolate evidence of the oblique effect. The understanding of this motion detection bias can be applied to more complex composite tactile stimuli used in real world environments, such as HMIs, to enhance user accuracy. The visual oblique effect, characterised similarly in isolated psychophysical experiments, has been reported in a variety of real-world contexts including product design and perception of fine art [Latto et al., 2000, Lidwell et al., 2010].

### Time-confidence trade off in complex tactile percepts

Remarkably, we found that accuracy was not affected by stimulus duration, despite the large range of durations used (Table S1). In contrast, confidence ratings were affected, which is a crucial additional consideration in the design of HMIs. Unexpectedly, the relationship between stimulus duration and confidence assumed a clear non-linear inverted U-shape (Figure 4): participants were least confident about the shortest and the longest durations. One explanation may be that longer exposure to the tactile percepts cause desensitisation, leading the perception to become less certain over time. This would decrease confidence, but not necessarily accuracy, if participants stick to their first choice. Our findings are somewhat unexpected, given that previous pilot studies using ultrasound stimuli have employed very long stimulus durations [Rutten et al., 2019]. Overall, our results suggest that long exposure to the tactile percepts is at the very minimum unnecessary, but also potentially detrimental to user experience. If this is an issue of desensitisation, in order to apply such ultrasound techniques to HMIs, there would be a clear advantage to selecting stimulus features that take advantage of perceptual biases (e.g. brief vertical motion) to enhance the accuracy of perceived feedback.

### Applications involving shapes

To our knowledge, this study is the first to rigorously test for the feasibility of whole hand tactile perception using ultrasound stimuli. To date, the majority of studies applying this technology have been usability pilots, which have focused on the parallels between ultrasound stimuli and a visual screen or display. As a result, most of this work has focused on shapes: a visual feature that appears intuitive to translate into the tactile domain. These pilot studies were proof-of-concept, and therefore employed limited sample sizes and/or small trials numbers, which preclude the use inferential statistics, limiting interpretability. However, this literature provides a foundation for the present study, and our desire to focus on motion rather than shape as a tactile feature for whole hand perception.

A recent example was a study testing fifteen participants in a shape-discrimination task Korres and Eid [2016]. On each trial, one of the four possible stimuli (line, circle, triangle, plus-sign) was presented for maximum 30 seconds, with the task consisting of 24 trials in total. Accuracy on the group level was highly variable across shapes (44-76%) - though these are difficult to interpret because each trial featured all stimuli as options. For example, the line stimulus was recognised correctly in 44% of trials, which is clearly above chance level, but it was misidentified as a circle in 51% of trials - which is concerning given the obvious spatial differences between lines and circles. Furthermore, reaction times (RT) were very slow (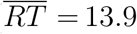 seconds over trials and participants). Before the experimental trials there was an unlimited period of training (times not reported), clouding interpretation of the results. Instead, Rutten et al. [2019] did not include any training, aiming to measure baseline performance. They tested a similar discrimination-task on 50 participants, with eight different stimuli (four static, four moving) that were presented for 9 seconds maximum per trial. The experiment consisted of 40 trials (5 blocks, each consisting of one trial for each stimulus). Accuracy was low to moderate (26-60% on group level across stimuli). It should be noted that their random-without-replacement design may produce progressive determination effects, meaning participants explicitly take their choice on trial *n-1* into account for their choice of trial *n*, making it difficult to determine chance level [Blais, 2008].

Other work has considered the application of virtual 3-D shapes using ultrasound: Long et al. [2014] asked participants to discriminate between five shapes (sphere, pyramid, horizontal prism, vertical prism, and cube), and found mean accuracy scores between 66.7% to 94.4% across shapes. Indeed, the exploring of edges seems more in line with the way we use our hands in daily life. Again however, power was low (6 participants with 15 trials each), and participants had an unlimited training period, necessitating further testing to definitively compare the perception of 2D vs 3D tactile shapes generated with ultrasound.

Although shape discrimination using active touch draws intuitive parallels between the tactile and visual system, this specific sensory feature may not be best suited for rapidly conveying sensory information via the hand. Shape discrimination relies on haptic exploration of a virtual object meaning motor behaviour accounts for participant variance. In contrast, motion stimuli targeted to the hand using infra-red tracking provide a greater degree of control over delivery, rendering them a more appealing mechanism for the rapid feedback required in touch-free interactions with HMIs [Breitschaft et al., 2019]. Importantly, tactile motion stimuli could also be integrated into touch interfaces that are not touch-free, for example, via actuators embedded in car steering wheels or clothing.

### Future Directions

While the question of feasibility and accuracy at the group level is important, the performance of *individual* participants in perceiving complex tactile percepts is relevant both from a mechanistic perspective (uncovering neurobiological processes) and from an applied perspective (testing feasibility for specific user-groups). We found large individual differences in performance that were not explained by our candidate variables. The lack of an observed relationship between performance and age likely resulted from a relatively young participant group. Tactile sensitivity in the fingertips is known to decrease with age and to be affected by gender, necessitating further evaluation of the accessibility of HMIs that result on ultrasound feedback [Goldreich and Kanics, 2003, Stevens et al., 1996, Thornbury and Mistretta, 1981, Goldreich and Kanics, 2003, Thornbury and Mistretta, 1981].

Another question of interest is to what extent whole hand tactile perception can improve with training. We found evidence against improvement in performance over time. However, aside from a few training trials, participants did not receive any feedback on their answers throughout. This could explain why some of our participants scored below chance level: they may have felt a difference between the two stimuli, but mislearned the association between stimulus and direction. Previous work on visual motion perception has found that training effects are usually limited. For example, training visual motion discrimination along a particular axis can improve performance, but this improvement does not carry over to performance along new axes [Ball and Sekuler, 1987]. This means that even if participants can learn whole hand motion discrimination with feedback, it is doubtful that this will show transfer effects to other tasks or even motion directions. The potential lack of a transfer effect will depend heavily on the end user. For example, in the context of users with sensory impairment, the prospect of prolonged training to learn individual stimulus types might be acceptable. In contrast, in the context of commercial HMIs in cars and clinical settings, such a learning curve would be less realistic. A more fruitful future approach may be to examine cross-rather than intra-modality training effects - training on visual and testing on tactile, or vice versa, to tap into the multisensory nature of perception.

## Conclusion

The current study is the first to investigate the perception of the whole hand complex tactile stimuli that have been made feasible with ultrasound techniques. In spite of the relatively sparse innervation of the palm compared with the fingertips, we found participants were able to perceive subtle moving dot stimuli above chance level. Using these stimuli were found clear evidence of an oblique effect in the perception of tactile motion across the hand. Motion aligned with the cardinal horizontal and vertical axes of the hand was perceived significantly more easily and confidently than that aligned with an oblique axis. In addition, participants felt most confident in the perception of stimuli around 500-2500 ms in duration.

A robust understanding for the perceptual biases in these complex tactile percepts will advance the implementation of touch-free tactile interfaces in practical contexts such as accessibility (e.g. haptic aids for visually impaired patients) and safety critical user interfaces (e.g. reducing visual overload in cars). The potential for mid-air tactile feedback to improve the accuracy of touch-free HMIs in clinical settings and busy public environments is also an attractive future application in the context of reducing the transmission of communicable diseases [Otter and French, 2009, Rossol et al., 2014]. However, such uses should avoid the temptation to directly translate stimuli, such as shape, from the visual domain into the tactile, albeit technically feasible. While we demonstrate that biases such as the oblique effect exist across sensory boundaries, vision and touch have unique predilections and acuities that, once identified, can be leveraged for practical purposes in HMIs.

## Acknowledgments

This work was funded by the MRC Proximity to Discovery grant (MC_PC_17186). JK is supported by a Sir Henry Wellcome Postdoctoral Fellowship ((204696 /Z/16/Z). The authors have no financial conflicts to disclose. We would like to thank Denny Han for his assistance with the testing lab setup, and UltraLeap Ltd for their technical support through the Ultrahaptics Academic Programme. Ultraleap saw the results of this manuscript ahead of publication. We are grateful to Brian Maniscalco for making their Matlab scripts used for the estimation of meta-cognitive measures publicly available. We also also grateful to Kristian Rusanov for his assistance with stimulus design.

## Supplementary Results

The Bayesian RM ANOVA on % correct with condition and duration as independent factors showed extremely strong evidence that participants’ accuracy was not dependent on the duration of the stimulus (BF_01_ = 14184) = see Supplementary Table S1 for all the means and standard deviations of accuracy over the different conditions.

**Supplementary Table S1.** Overview of the mean (SD) of % correct over the different durations, for each of the three conditions and for all conditions combined

| Duration | Combined | Vertical | Horizontal | Oblique |
| --- | --- | --- | --- | --- |
| D1 | .60 (.11) | .62 (.14) | .60 (.14) | .57 (.13) |
| D2 | .61 (.13) | .61 (.17) | .61 (.15) | .60 (.16) |
| D3 | .62 (.13) | .62 (.19) | .62 (.15) | .61 (.16) |
| D4 | .60 (.16) | .60 (.19) | .61 (.18) | .60 (.16) |
| D5 | .61 (.13) | .64 (.15) | .61 (.16) | .58 (.17) |
| D6 | .60 (.15) | .61 (.18) | .60 (.18) | .59 (.16) |
| D7 | .61 (.18) | .63 (.20) | .64 (.21) | .58 (.19) |
| D8 | .60 (.16) | .62 (.21) | .61 (.18) | .56 (.18) |
| D9 | .61 (.17) | .62 (.20) | .63 (.20) | .58 (.20) |
| D10 | .61 (.17) | .65 (.19) | .62 (.21) | .57 (.21) |

